# Near-perfect discrimination of half-sibling and avuncular pairs using genomic data without pedigrees

**DOI:** 10.64898/2026.01.06.697070

**Authors:** Emmanuel Sapin, Kristen M. Kelly, Matthew C. Keller

## Abstract

**Motivation:** Large-scale genomic biobanks contain thousands of second-degree relatives without pedigree data. Accurately distinguishing half-sibling and avuncular pairs–both of which share approximately 25% of the genome–remains a significant challenge. Current SNP-based methods rely on aggregate Identical-By-Descent (IBD) segment counts and age differences, but substantial overlap in these distributions leads to high misclassification rates. There is a need for a scalable, genotype-only method that can resolve these second-degree ambiguities without requiring observed pedigrees or phenotypic information.

**Results:** We present a novel computational approach that achieves near complete separation of half-siblings and avuncular pairs using haplotype-level sharing features based on data that has been computationally phased across-chromosomes. By modeling these features with a multivariate Gaussian mixture model, we achieve exceptional classification performance in biobank-scale data. Ground-truth labels for validation were established through a multi-step inference process within a family graph constructed from high-confidence first, second and third-degree relationships. All 444 ground-truth avuncular pairs and 97 of the 99 half-sibling pairs were correctly classified. Treating avuncular status as the positive class, this produces a specificity of 97.98% and sensitivity of 100%. These features also make it possible to determine which member of an avuncular pair is the niece/nephew and which is the avuncular with 99.77% empirical accuracy (443/444 reference pairs correctly polarized). Our results suggest that the framework could perform comparably on other similar datasets. This method provides a robust, scalable means of distinguishing half-sibling and avuncular pairs, improving relationship annotation for pedigree inference, genomic analyses that utilize relatives, and forensic genetic applications.

**Contact:**, and

## 1 Introduction

Genome-wide datasets routinely contain closely related individuals without information about pedigree relationships, which must be inferred from genotype data (10). As sample sizes in biobank-scale cohorts have grown to hundreds of thousands of individuals, accurately inferring these relationships has become increasingly important for reconstructing pedigrees, improving haplotype phasing, and supporting downstream analyses in human genetics, including demographic inference and genealogical applications (2; 15; 4).

A persistent challenge in relatedness inference is distinguishing between relationships beyond the first degree. Parent–offspring and full sibling pairs can be readily distinguished because, although both share approximately 50% of their genome identical-by-descent (IBD), parent–offspring pairs share no genomic segments where both alleles are inherited from the same ancestor (IBD2), whereas full siblings share 25% of their genomes IBD2. While theoretical inbreeding configurations such as double grandparent–grandchild pairs (resulting from offspring between two half-siblings) would share the same sharing pattern as full siblings (with both IBD1 and IBD2 sharing), such high-inbreeding structures are vanishingly rare in outbred human populations, are routinely excluded from human genetic datasets, and are consequently not considered here. This makes standard first-degree relationships simple to classify perfectly from genomic data. In contrast, several distinct pedigree configurations produce the same expected sharing for second-degree relatives (an average of 25% IBD), including half-siblings, grandparent–grandchild, avuncular (niece/nephew–uncle/aunt), and double first-cousin pairs (pairs in which both parents of one cousin are siblings of both parents of the other cousin), and the genome-wide patterns of IBD sharing are similar across these relationship classes (13; 10). Among these, half-sibling and avuncular pairs are the most important to distinguish in biobank data because they are by far the most common. Grandparent–grandchild pairs are rare in cohorts recruited from a single adult generation, and double first cousin pairs are exceedingly rare in most samples.

Accurately distinguishing half-sibling from avuncular relationships is important across several domains of genomic research. Because the two relationships imply different pedigree structures, distinguishing them improves pedigree reconstruction in large cohorts and analyses that depend on correctly specified family relationships. The distinction may also affect quantitative genetic analyses: half-siblings are more likely than avuncular pairs to share childhood environments and direct parental influences, so misclassification can bias estimates derived from pedigree data (16). In forensic genetics, correct classification can likewise be essential because lineage inference and individual identification depend on reconstructing the correct family configuration (13; 7).

Current approaches to distinguish half-sibling from avuncular pairs primarily rely on the distribution of IBD segments; half-siblings inherit shared genetic segments directly from their common parent, whereas segments shared by niece/nephew-avuncular pairs have undergone an additional meiosis, typically leading to a higher count of shorter segments (2). Empirical distributions of segment length and count nonetheless overlap substantially, a consequence of the stochastic nature of recombination (14). Pairwise classifiers built on these aggregate diploid statistics correctly resolve only 92.8–98.6% of second-degree pairs as a class, without separating half-sibling from avuncular performance (10).

Haplotype-based scores improve on this baseline by modeling background sharing in endogamous or structured populations, but require training on known relative pairs from the same sample (14). In two independently validated cohorts, this approach achieves 82.7–96.8% sensitivity for avuncular pairs and 85.5–94.7% for half-siblings (14). The dependence on in-sample training data limits the approach to cohorts with enough known relative pairs to train on, a condition many large, sparsely related biobank samples do not meet.

A separate class of methods instead draws on genotyped collateral relatives shared by the pair, inferring relationship type from multi-way IBD sharing across additional family members (12; 9). In real pedigree data from Generation Scotland, CREST achieves 97.7% sensitivity for avuncular pairs and 92.2% for half-sibling pairs with sufficient mutual relative data (9). PADRE, an earlier composite-likelihood method in this class, was evaluated only in simulation, where its avuncular sensitivity ranges from 60.5% to 92.0% across seven pedigree structures, falling lowest when the only available mutual relative is a second cousin rather than a closer collateral relative (12; 9). Most second-degree pairs identified in biobank-scale samples lack the genotyped collateral relatives these methods require, restricting their use to the minority of pairs embedded in larger genotyped pedigrees. Demographic information such as age differences can provide additional heuristic support in these cases, but such data are not always available and are rarely sufficient on their own for high-confidence classification.

Each of these approaches has different limitations. Diploid IBD statistics use only the pair’s own genotypes, but discard haplotype structure and are not very accurate at separating half-sibling from avuncular pairs. Haplotype-score and multi-way IBD methods recover this structure and classify more accurately, but neither is close to perfect, and both require additional known relatives beyond the pair – other relative pairs to train on, or genotyped collateral relatives in the pedigree. Biobank cohorts, sampled from distantly related individuals, often supply neither. Most second-degree pairs have no additional genotyped relatives in the sample, nor is there a cohort of known relationships large enough to train a classifier.

In this paper, we propose a scalable, genotype-only computational framework for distinguishing the two most common types of seconddegree relationships in biobank data without requiring explicit pedigree or phenotypic information. As a first step, we apply a recently developed computational algorithm to assign haplotypes to consistent parental origins across chromosomes (11). We then summarize the across-chromosome distribution of IBD sharing in a two-dimensional feature vector that captures the distinct inheritance patterns of half-sibling and avuncular pairs. In half-siblings, all shared IBD segments derive from the same parent, whereas in avuncular pairs, shared segments are confined to one haplotype of the niece or nephew but may lie on either haplotype of the aunt or uncle. Modeling these features with a multivariate Gaussian mixture model fitted using the Expectation-Maximization algorithm (5) provides near-perfect separation between the two relationships despite unavoidable errors in computationally inferred across-chromosome phase. Once classified, these relationships can also inform across-chromosome phasing: half-sibling pairs provide particularly informative phase anchors because all shared IBD segments identify the same parental lineage, whereas avuncular pairs provide a less direct constraint. This enables a feedback loop that can further improve the performance of across-chromosome phasing.

## 2 Methods

### 2.1 UK Biobank sample and ancestry definition

The method was developed and evaluated using data from the UK Biobank (3), focusing on individuals of European descent. This choice was motivated by the demographic composition of the UK Biobank cohort, where non-European ancestry groups have substantially smaller sample sizes that currently limit the accuracy of across-chromosome phase estimation.

Genetic principal components (PCs) were computed on a linkage-disequilibrium–pruned set of common autosomal SNPs. The smallest 4-dimensional hypercube in PC space containing all individuals with self-described white British ancestry was then identified, and all individuals lying within this hypercube, regardless of self-described ancestry, were retained as the eligible sample (435,187 individuals).

Next, we selected biallelic autosomal SNPs that were present on both the UK BiLEVE and UK Biobank Axiom arrays, had a minor allele frequency (MAF) of at least 5%, and had a Hardy–Weinberg equilibrium chi-square test *p*-value greater than 0.0005. This resulted in a final dataset of 327,744 SNPs for relatedness estimation and downstream analyses (3).

### 2.2 Detection of 2nd-degree relatives

We calculated the standard measures of diploid SNP similarity (8), 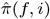and 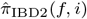, as defined in Equations 1 and 2.

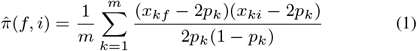

In this equation, *f* denotes the focal individual, *i* indexes the non-focal individuals, *p*_*k*_ is the reference allele frequency at SNP *k, x*_*kf*_ and *x*_*ki*_ represent the diploid genotypes (coded as 0, 1, or 2 copies of the reference allele) for individuals *f* and *i* at SNP *k*, and *m* is the total number of SNPs included in the calculation.

To determine the genome-wide proportion of genomic segments where individuals share both homologous alleles IBD at a locus (IBD2), we calculated 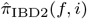 as the average standardized product of the dominance deviations for the focal individual *f* and non-focal individual *i* across SNPs, as shown in Equation 2.

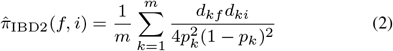

This follows the standard method-of-moments framework implemented within the PLINK software toolkit (8), where *d*_*kf*_ represents the dominance deviation for the focal individual *f* at SNP *k* (and similarly *d*_*ki*_ for individual *i*), defined conditionally based on the diploid genotype *x*_*kf*_ *∈ {*0, 1, 2*}*:

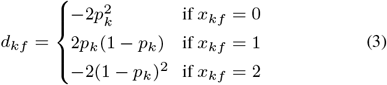

All UK Biobank participants were aged 40–69 at the time of recruitment (3). Given that a grandparent–grandchild (GP–GC) pair requires two successive generational transitions, the minimum demographic age gap between a grandparent and grandchild is typically at least 30–35 years. Within the narrow 29-year recruitment window (ages 40–69), a GP–GC relationship would necessitate an extreme reproductive age distribution (e.g., both parent and grandparent reproducing in their mid-teens, combined with recruitment at the extreme boundaries of 40 and 69 years). Consequently, true GP–GC pairs are expected to be virtually absent.

Nevertheless, because our Gaussian Mixture Model (GMM) partitions the feature space into two primary classes (half-siblings vs. avuncular pairs), we explicitly evaluated the risk of contamination from remaining alternative second-degree or anomalous configurations (such as residual GP–GC pairs, double first cousins, or pairs with cryptic endogamy). Double first cousins share on average 25% of their genome 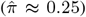 but are characterized by non-zero IBD2 sharing 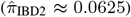, whereas true half-sibling and avuncular pairs exhibit 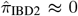. By applying an initial filter requiring 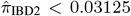 (half of the two 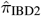 values 0 and 0.0625), double first cousins and pairs reflecting close inbreeding were systematically removed. Furthermore, theoretical haplotype transmission dictates that GP–GC pairs share segments derived from only one parental gamete across all chromosomes, yielding across-chromosome phasing and sharing signatures that mirror the uniparental distribution of avuncular pairs rather than half-siblings. Thus, while any undetected GP–GC pair would coalesce into the avuncular cluster rather than inducing a spurious third cluster, the extreme recruitment age window and kinship filtering ensure that the 8,728 candidate pairs 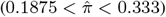 are essentially composed of half-sibling and avuncular relationships.

### 2.3 Ground truth labeling

To obtain labeled pairs of specific 2nd-degree types, a family graph was constructed from high-confidence 1st-degree relationships. Parent–child and full-sibling relations were identified using 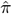 and 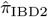 sharing. From these relationships, the following labels could be assigned to a subset of our identified second-degree relatives:

- **Half-siblings**: a pair of individuals sharing one parent.
- **Avuncular pairs**: a pair in which one individual is a child of the other individual’s sibling.

This procedure led to the identification of 444 avuncular pairs but only 3 half-sibling pairs. To increase the number of verified half-sibling pairs, we used information from other degrees of relatedness, despite the additional uncertainty. Two configurations involving third-degree relationships (with 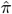 around 0.125) can be used to identify half-siblings with high confidence, leading to a set of pairs with high-confidence pedigree-inferred (HCPI validation set) relationships that we use as a reference to assess the method. While we frame these configurations in terms of first cousins, they could alternatively involve half-avuncular pairs (i.e., a half-sibling of a parent), as both relationship types share the same expected relatedness 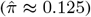 and follow the exact same lineage segregation logic detailed below:

- **Reciprocally unrelated first cousins**: pairs in which each individual (*half* -*sib*_1_ and *half* -*sib*_2_) has a first cousin (*FC*_1_ and *FC*_2_) who is unrelated to the other member of the pair (Figure 1). This configuration is compatible with half-siblings but is incompatible with an avuncular relationship. In an avuncular pair (*A, N* ), a first cousin of the niece or nephew could be unrelated to the aunt or uncle if the relationship occurs through the niece or nephew’s other parent. However, any first cousin of *A* would also be a first cousin of *N* ‘s parent and therefore a first cousin once removed of *N*, with expected relatedness 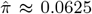 (Figure 2). Thus, the aunt or uncle cannot have a first cousin who is unrelated to the niece or nephew, and pairs exhibiting the reciprocal pattern were classified as half-siblings.
- **Shared first cousin**: pairs in which the two individuals (*half* -*sib*_1_ and *half* -*sib*_2_) share a first cousin (*FC*) must be half-siblings, as illustrated in Figure 3. Such pairs cannot be avuncular, because, as illustrated in Figure 2, an avuncular individual’s first cousin (*CA*) would share 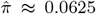 with *N* rather than *≈* .125. Therefore, these pairs are classified as half-sibling.

**Figure 1.**
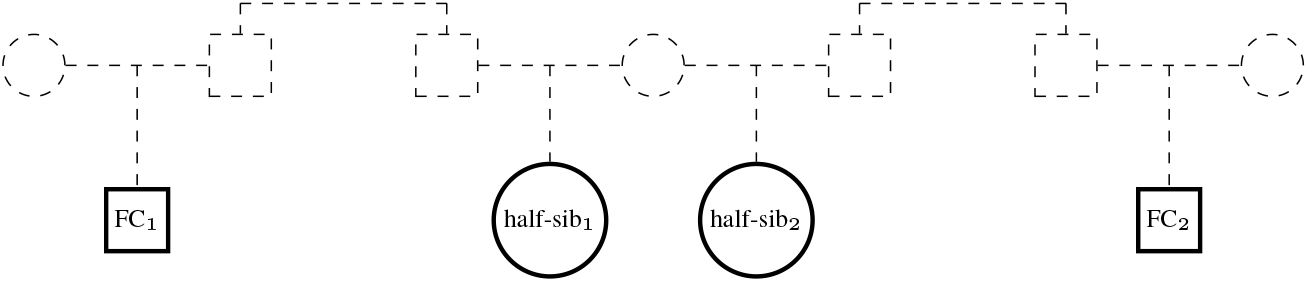
Example pedigree showing first cousins of a half-sibling pair. The half-siblings share a mother, and each has a paternal first cousin (FC_1_, FC_2_). Each half-sibling is unrelated to the other’s first cousin, a configuration that would be impossible for an avuncular pair.

**Figure 2.**
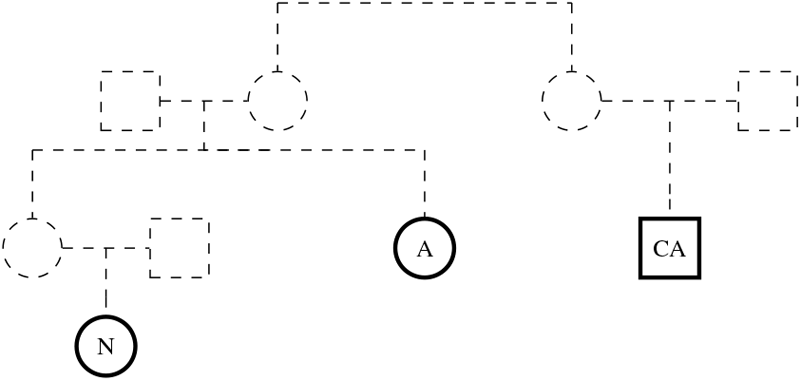
Example pedigree showing a first cousin of an avuncular relative. The first cousin *CA* of the avuncular relative *A* is also the first cousin of *A*’s sibling (the parent of *N* ). Consequently, in an avuncular pair, *CA* and *N* (the niece/nephew) must be related (expected 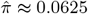, first cousin once removed).

**Figure 3.**
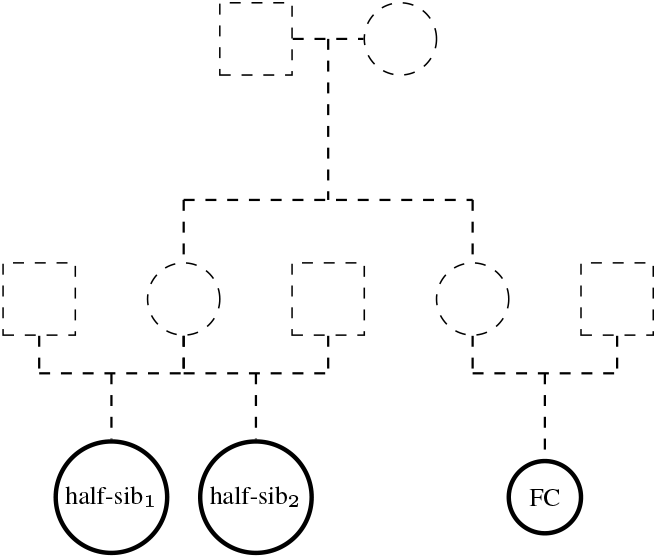
Example pedigree showing two half-siblings (*half* -*sib*_1_, *half* - *sib*_2_) sharing a mother and a maternal first cousin (*FC*). The first cousin has the same degree of relatedness to both half-siblings, which would be not be the case for an avuncular pair.

This procedure successfully recovered 99 half-sibling pairs, which were incorporated alongside the 444 avuncular pairs into the High-Confidence Pedigree-Inferred (HCPI) validation set to serve as ground truth for evaluating our classification framework.

The remaining second-degree pairs that could not be labeled as half-siblings with high confidence—due to their lack of informative relatives within the sample or due to cousin relationships potentially consistent with an avuncular relationship—were left unlabeled. When establishing the HCPI validation set for half-sibling detection, relatives were classified as first cousins using the conservative threshold of 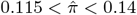 .Pairs following a scenario in which a cousin shares ancestry with both of an individual’s parents need to be excluded from the HCPI reference set (remaining in downstream analyses as unlabeled candidate pairs). Those scenarios would result in 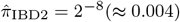 between an individual in the pair and their cousin as a consequence of independent Mendelian sampling across both paths (6), which could distort the analysis of standard outbred relationships. A cutoff of half this value was therefore chosen, excluding pairs from the HCPI validation set for which 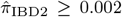 between an individual in the pair and their cousin. Finally, a cousin of one individual was considered unrelated to the other individual in the pair if 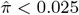. Pairs excluded from the HCPI set of pairs were still included in the later analysis of unlabeled pairs.

### 2.4 Across-chromosome phasing

All UK Biobank individuals were initially phased within chromosomes using Shapeit2 (1; 3). To extend phase across chromosomes, we applied the across-chromosome phasing method described in (11). This method assigns chromosome-specific haplotypes to a consistent parental origin, thereby identifying which haplotypes across different chromosomes were inherited from the same parent, by leveraging genome-wide patterns of haplotype sharing.

The resulting across-chromosome haplotype assignments are probabilistic rather than deterministic. In validation analyses using UK Biobank offspring with both parents genotyped as a ground truth, but with parental genotypes withheld during phasing, the method achieved a mean across-chromosome phasing accuracy of 83.1% when applied to Shapeit2-phased data (median = 85.9%). When within-chromosome phase errors were removed, mean accuracy increased to 95.3% (median = 100%), indicating that most of the error arises from imperfect within-chromosome phase rather than from the across-chromosome phasing procedure itself (11).

For the ground-truth pairs in this analysis, we applied the same across-chromosome phasing restriction used by Sapin et al. (11): parents, other first-degree relatives, and second degree relatives of each focal individual were excluded from the reference set, while more distant relatives were retained. This prevents the ground-truth pairs from being phased with directly informative close relative data, which would inflate phasing accuracy and, in turn, overestimate the sensitivity and specificity of the half-sibling versus avuncular classifier. The use of more distant relatives is appropriate, because such sharing is part of the information the algorithm uses and reflects the setting in which candidate second-degree relationships are classified.

The central idea of across-chromosome phasing is that, although chromosomes are initially phased independently, haplotypes inherited from the same parent will exhibit systematically higher genetic similarity to the same set of external individuals (i.e., distant relatives) across the genome. This occurs because a common ancestor of the focal and non-focal individuals may transmit IBD segments on multiple chromosomes, and—absent inbreeding—these segments must be inherited through the same parental lineage in the focal individual. As a result, haplotypes from that parent will show elevated SNP similarity to the same non-focal individuals across multiple windows on different chromosomes, producing correlated similarity profiles, whereas haplotypes from different parents will not.

To capture this signal, the genome is partitioned into windows defined by recombination hotspots. Within each window, each haplotype of a focal individual is compared to both haplotypes of all non-focal individuals in a reference sample. For each comparison, we retain the maximum SNP similarity, which reduces sensitivity to phase errors in the reference panel. This produces, for each focal haplotype and each genomic window, a vector of similarity scores to the reference individuals.

Across windows and across chromosomes, haplotypes originating from the same parent will show correlated patterns of similarity to the reference sample, whereas haplotypes from different parents will not. The algorithm therefore computes correlations of these similarity profiles across windows and iteratively groups haplotypes into two clusters corresponding to the two parental homologues (*p*_1_ and *p*_2_). By aggregating information across the genome in this manner, the method establishes a consistent parental assignment across all autosomes without requiring pedigree information or explicit IBD segment detection.

### 2.5 Haplotype-level sharing metrics for relationship classification

For each candidate second-degree pair, we computed four genome-wide haplotype–haplotype similarity values, denoted 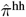. These correspond to the four pairwise comparisons between the two across-chromosome phased haplotypes of individual *i* and the two across-chromosome phased haplotypes of individual *j*. The standard diploid statistic 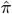 in Equation 1 measures similarity between diploid genotypes. Here, we decompose that statistic into haplotype-level components.

For haplotype *a* of individual *i* and haplotype *b* of individual *j*, with *a, b ∈ {*1, 2*}*, we define

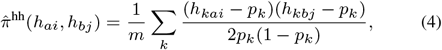

where *h*_*kai*_ and *h*_*kbj*_ denote haploid genotypes at SNP *k*, coded as 0 or 1 copies of the reference allele, and *p*_*k*_ is the reference allele frequency. The factor of 2 in the denominator places the haplotype–haplotype values on the same scale as the diploid statistic: the four 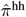 values sum to the diploid 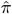 for the pair. Thus, a haplotype pair sharing proportion *s* of the genome IBD has expected 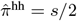.

This scaling gives simple expectations for the two relationship classes. For half-siblings, the two haplotypes inherited from the shared parent are expected to share half their genome IBD, yielding one elevated value with 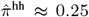. The other three haplotype–haplotype comparisons are expected to be near zero. For avuncular pairs, IBD segments transmitted to the niece/nephew pass through the intervening parent and therefore reside on one haplotype of the niece/nephew. In the avuncular, the corresponding shared segments may lie on either haplotype inherited from the shared grandparents. Thus, two of the four 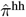 values are expected to be approximately 0.125, and the remaining two are expected to be near zero, as illustrated in Figure 4.

**Figure 4.**
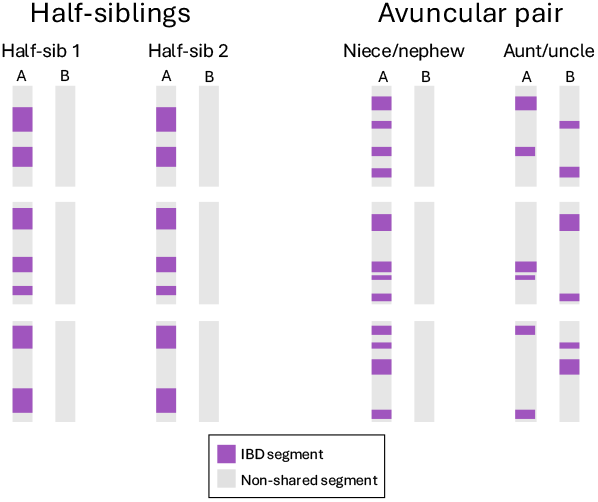
Example distribution of IBD sharing across across-chromosome phased haplotypes for half-sibling and avuncular pairs. In half-siblings, shared segments derive from one shared parent and are concentrated in one haplotype–haplotype comparison. In avuncular pairs, shared segments are confined to one haplotype of the niece/nephew but distributed across both haplotypes of the avuncular.

The four values form a 2 *×* 2 matrix for each pair (Table 1). Because haplotypes are unlabeled with respect to parental origin, we relabel rows and columns post hoc using only the four observed 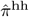 values, without using relationship labels. We order the matrix so that 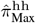 is the largest entry. Among the two entries in the same row or column as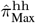, we label the larger value 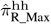 and the smaller value 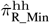 . The remaining entry, diagonally opposite 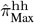, is denoted 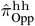.

**Table 1.**
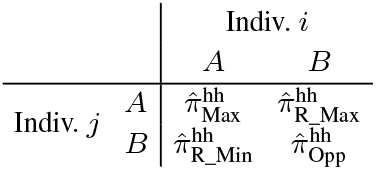
Haplotype–haplotype 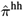 values for a pair of individuals. Haplotypes are indexed as *A, B* for individuals *i* and *j*, and are relabeled post hoc so that 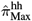 is the largest entry.

|  |  | Indiv. *i* |  |
|---|---|---|---|
|  |  | *A* | *B* |
| Indiv. *j* | *A* | π̂<sup>hh</sup><sub>Max</sub> | π̂<sup>hh</sup><sub>R_Max</sub> |
|  | *B* | π̂<sup>hh</sup><sub>R_Min</sub> | π̂<sup>hh</sup><sub>Opp</sub> |

The genome-wide 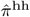 values above describe the expected inheritance patterns under ideal phasing, but they are not sufficient on their own for classification in real data. Across-chromosome phasing is imperfect, and phase errors add noise to the 2 *×* 2 haplotype-sharing matrix. We therefore partitioned SNPs by genotype status to construct metrics that accentuate the phase-specific differences between half-sibling and avuncular pairs. Each of the two individuals in the pair was classified at each SNP as either homozygous (Hom) or heterozygous (Het), producing four possible genotype-pair categories (Hom-Hom, Hom-Het, Het-Hom, and Het-Het). Only two genotype-pair classes were used in the final metrics. HetHet SNPs are phase-informative in both individuals and provide the clearest relationship-specific signal. Double-homozygous SNPs are not phase-informative, because each individual carries the same allele on both haplotypes. They nevertheless provide a phase-independent baseline for overall SNP similarity: matching homozygotes contribute positively to all four haplotype–haplotype comparisons, whereas discordant homozygotes contribute negatively. We use this baseline below to scale the Het-Het metrics, so that the classifier emphasizes phase-specific Het-Het sharing relative to each pair’s background SNP similarity, rather than differences in overall pairwise similarity. Mixed homozygote–heterozygote categories were excluded because they are phase-informative for only one individual and therefore cannot identify a specific haplotype–haplotype pairing.

At doubly heterozygous SNPs within IBD segments, half-siblings and avuncular pairs generate different haplotype-level patterns. Table 2 gives the contribution of a doubly heterozygous SNP *k* to each of the four haplotype–haplotype sums in Equation 4.

**Table 2.**
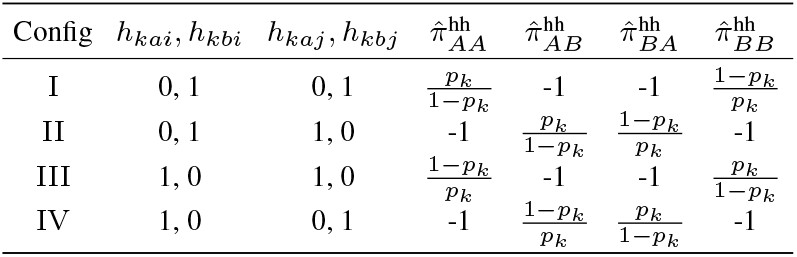
Contribution of SNP *k* to the pairwise similarity metrics for the four configurations of a doubly heterozygous SNP. We ordered the haplotypes for the two individuals following the conventions in Table 1. Note that while 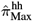 represents the genome-wide maximum similarity corresponding to respective *A* haplotypes of individuals *i* and *j*, the specific 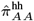 contribution at SNP *k* to this metric is not necessarily the maximum of the four 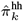 contributions at SNP *k*.

| Config | *h*<sub>*kai*</sub>, *h*<sub>*kbi*</sub> | *h*<sub>*kaj*</sub>, *h*<sub>*kbj*</sub> | *π̂*<sup>hh</sup><sub>*AA*</sub> | *π̂*<sup>hh</sup><sub>*AB*</sub> | *π̂*<sup>hh</sup><sub>*BA*</sub> | *π̂*<sup>hh</sup><sub>*BB*</sub> |
|---|---|---|---|---|---|---|
| I | 0, 1 | 0, 1 | $\frac{p_k}{1-p_k}$ | -1 | -1 | $\frac{1-p_k}{p_k}$ |
| II | 0, 1 | 1, 0 | -1 | $\frac{p_k}{1-p_k}$ | $\frac{1-p_k}{p_k}$ | -1 |
| III | 1, 0 | 1, 0 | $\frac{1-p_k}{p_k}$ | -1 | -1 | $\frac{p_k}{1-p_k}$ |
| IV | 1, 0 | 0, 1 | -1 | $\frac{1-p_k}{p_k}$ | $\frac{p_k}{1-p_k}$ | -1 |

For half-siblings, after assigning the shared-parent haplotypes the same label in both individuals, doubly heterozygous SNPs within IBD segments are expected to occur predominantly in configurations I or III (and only in these configurations when phase is perfect). At these SNPs, the two aligned haplotype comparisons, 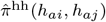 and 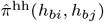, receive positive contributions, whereas the two cross-haplotype comparisons receive negative contributions. Because half-siblings share approximately 25% of their genome IBD on average, aggregating these negative contributions across shared segments can lead to negative empirical values for the two cross-haplotype metrics 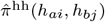 and 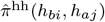. Doubly heterozygous SNPs outside IBD segments do not exhibit this systematic alignment and contribute background variation across the four configurations. Consequently, when contributions are aggregated across the genome, half-siblings exhibit stronger similarity in the aligned haplotype comparisons than in the cross-haplotype comparisons.

For avuncular pairs, the shared allele lies on one haplotype of the niece/nephew but can align with either haplotype of the avuncular. Across doubly heterozygous SNPs, this makes the four configurations I, II, III, and IV approximately balanced, so the expected Het-Het contribution to each of the four 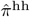 values is near zero. This conditional expectation differs from the all-SNP expectation because the genome-wide 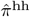 values also include homozygous and other non-Het-Het sharing. Restricting to Het-Het SNPs accentuates the phase-specific contrast between the two relationship classes, which helps compensate for noise introduced by imperfect across-chromosome phasing.

We evaluated the haplotype–haplotype entries in Table 1 separately for doubly homozygous and doubly heterozygous SNPs. At a doubly homozygous SNP, the four SNP-level haplotype–haplotype contributions are identical because, within each individual, the two haplotypes carry the same allele; the two individuals may nevertheless have matching or discordant homozygous genotypes. We aggregate these contributions across all doubly homozygous SNPs to obtain 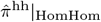, a phase-independent baseline for overall SNP similarity. For doubly heterozygous SNPs, we retain the matrix ordering defined in Table 1 and aggregate contributions across the Het-Het subset. We denote the largest genome-wide Het-Het entry as 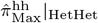 and the smallest of the remaining three entries as 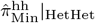 . These two Het-Het values are then scaled by 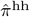 in Equations 5 and 6.

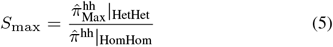

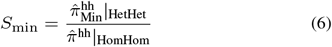

The resulting two-dimensional haplotype-sharing vector for each candidate second-degree pair is

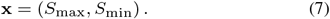

### 2.6 Gaussian mixture model and posterior probability *P* (half-sibling)

To classify candidate second-degree pairs, we model the two-dimensional haplotype-sharing vector *x* = (*S*_max_, *S*_min_) as a mixture of two equal-covariance Gaussians, one for half-siblings and one for avuncular pairs. We fit a two-component Gaussian mixture with shared covariance by EM on HCPI pairs, initializing the means at the class centroids. Features are first centered and scaled using all complete pairs. Unlabeled pairs do not contribute to the fit.

The marginal density of *x* is

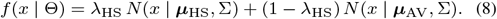

For any pair, the posterior probability of the half-sibling class is

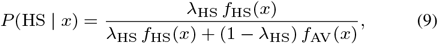

where *f*_HS_(*x*) = *ϕ*(*x* | ***µ***_HS_, Σ) and *f*_AV_(*x*) = *ϕ*(*x* | ***µ***_AV_, Σ) are bivariate normal densities. We call a pair half-sibling if *P* (HS | *x*) *>* 0.5 and avuncular otherwise. Unlabeled candidate pairs are scored with this same posterior; they do not contribute labels to the fit. How labeled HCPI pairs are trained and scored without using a pair in its own parameter estimates is described in the next subsection.

### 2.7 Cross-validation and runtime

Labeled HCPI pairs were scored with stratified 5-fold cross-validation.In each fold, EM was run on the training HCPI pairs only. Each labeled pair received the out-of-fold posterior P(HS). Unlabeled pairs were scored as the mean of the five fold models. Each labeled pair received the out-of-fold posterior *P* (HS) from the fold that held it out. Unlabeled pairs were scored as the mean of the five fold-wise linear discriminant analyses. On *n* = 8601 candidate pairs, the classification path took approximately 30 s to read the input, 15 s to apply the exclusion and 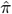 cutoffs, 20 s to fit the classifier, 1 s to score unlabeled pairs, and 30 s to write the prediction table (*≈* 96 s in total).

## 3 Results

### 3.1 Supervised cluster identification and validation

We classified second-degree relatives using the 5-fold EM mixture described above. The 2-D multivariate GMM demonstrated near-perfect classification performance when applied to the UK Biobank candidate set, identifying two clearly separated clusters within the HCPI reference set based solely on the 2-D feature vector (Figure 5). Using a decision boundary of *P* (half-sibling) *>* 0.5, all 444 reference avuncular pairs and 97 of the 99 reference half-sibling pairs were correctly assigned, yielding a sensitivity of 100% and a specificity of 97.98% (treating avuncular pairs as the positive class).

**Figure 5.**
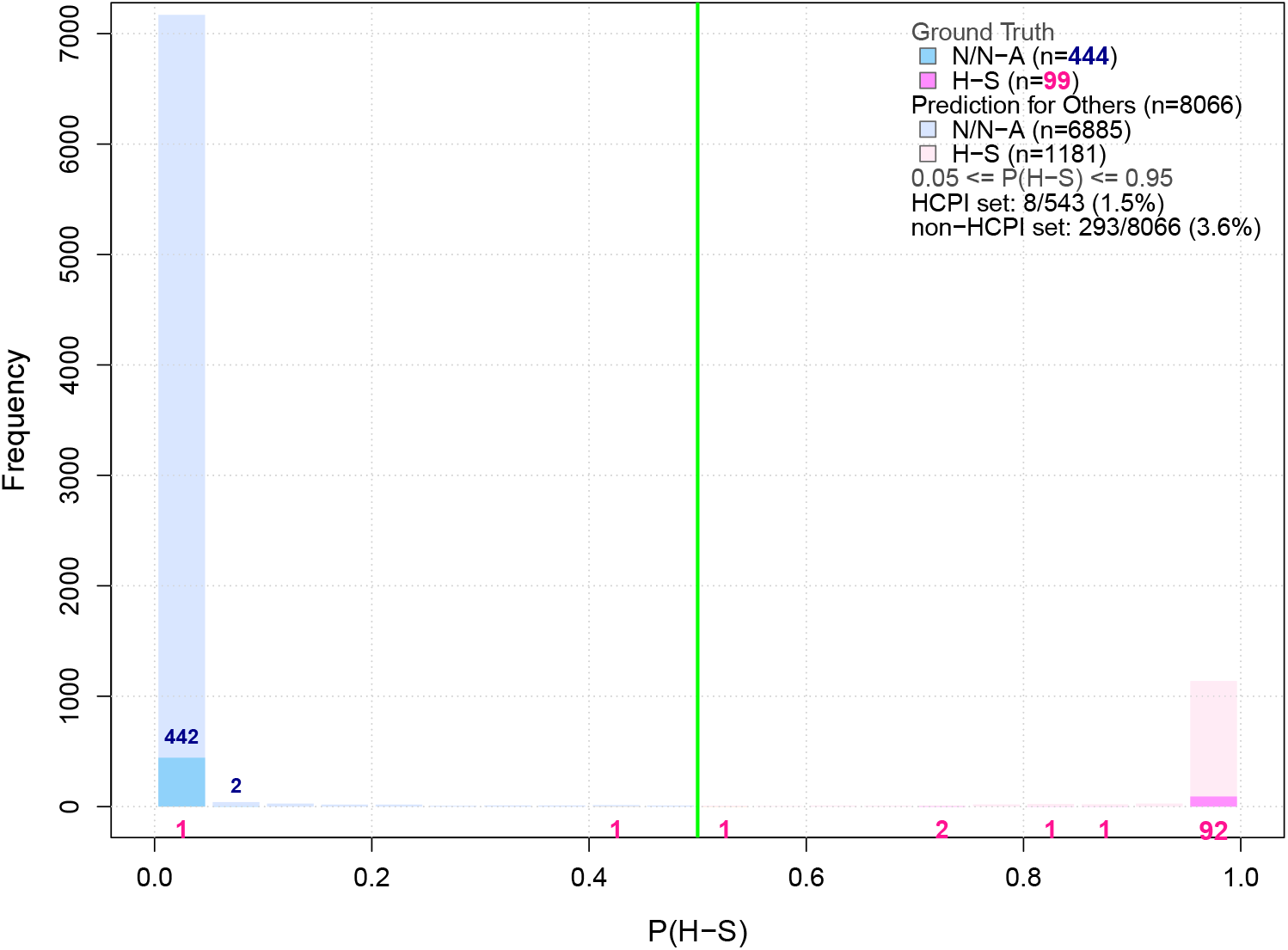
Distribution of the posterior probability *P* (half-sibling) for HCPI set of pairs and unlabeled second-degree relative candidates. The green line indicates the threshold for half-sibling identification (*P* (half-sibling) *>* 0.5). Blue numbers indicate number of HCPI avuncular relatives and pink numbers indicate the number of HCPI half-siblings.

While these validation metrics confirm exceptional performance in the UK Biobank cohort, generalizing these empirical results to external biobanks cannot be assumed solely from matching total sample sizes. Because the classification framework directly relies on across-chromosome phasing accuracy, its portability is fundamentally anchored by the density and distribution of informative distant relatives that scaffold parent-of-origin assignments. In cohorts of comparable size, differing demographic histories, higher structural heterogeneity, fine-scale stratification, or pervasive endogamy could depress background segment sharing or diminish the baseline quality of within-chromosome phase. Consequently, populations with sparser distant kindred networks may experience lower across-chromosome phasing fidelity, which would widen cluster variance and attenuate the separation observed between half-sibling and avuncular distributions. Demonstrating high empirical accuracy on our reference set therefore establishes proof-of-concept feasibility under favorable genealogical density, but formal validation across ancestrally diverse biobanks will be necessary to determine the precise kindred thresholds and marker densities required to replicate these separation boundaries.

### 3.2 Classification of unlabeled pairs and ascertainment

Beyond the HCPI reference set, the classification framework was applied to evaluate the remaining 8,066 previously unlabeled second-degree relative pairs in the UK Biobank cohort. The model classified 1,181 pairs as likely half-siblings (*P* (HS) *>* 0.5) and 6,885 pairs as likely avuncular pairs (*P* (HS) *≤* 0.5).

Because the HCPI reference set required specific genealogical configurations (e.g., reconstructed first-degree links for avuncular pairs and informative first-cousin configurations for half-siblings), pairs in the reference set are not an unconditional random sample of all second-degree relatives in the cohort. Specifically, we assessed whether denser local pedigree sampling in the reference set artificially enhanced within- or across-chromosome phasing accuracy:

- **Avuncular reference pairs:** Ascertained via a reconstructed first-degree link (e.g., an individual *T* who is the avuncular relative’s sibling and the niece/nephew’s parent). Although *T* was explicitly excluded from both within-chromosome phasing (3) and across-chromosome phasing (11), their presence could reflect broader kindred representation.
- **Half-sibling reference pairs:** Ascertained through direct common parentage, a shared first cousin, or reciprocal unshared first cousins. Additional third-degree relatives could theoretically provide subtle scaffolding during population-based phasing.

To quantify familial context in the unlabeled cohort relative to the labeled cohort, we evaluated empirical relatedness 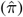 to each pair’s closest external relative (excluding first- and second-degree relatives; Supplementary Figure S4). The mean external 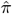 in the unlabeled cohort (0.0838) aligns closely with reference avuncular pairs (0.0846), whereas reference half-siblings exhibited higher values 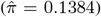. To evaluate the extent to which this difference impacts classification, we assessed the sharpness of the bimodal separation. Both cohorts demonstrated sharp bimodal separation toward the mixture components, with only 3.6% of unlabeled pairs (293 of 8, 053) and 1.5% of reference pairs (8*/*543) falling in the intermediate interval 0.05 *≤ P* (HS) *≤* 0.95 (Supplementary Figure S2). This close agreement (3.6% versus 1.5%) reflects consistent clustering stability, with the minor difference attributable to modest contamination in the uncurated unlabeled sample from rare, unmodeled configurations (such as residual grandparent–grandchild pairs or complex compound degree-2 relationships) that were excluded from the reference set by genealogical triangulation. Predictive performance is empirically substantiated by 5-fold cross-validation on the HCPI reference set (98.53% accuracy), confirming that the framework reliably separates genealogically isolated pairs, even when lacking the extended familial networks observed in reference half-siblings.

### 3.3 Misclassification of Two Pairs

Among the HCPI reference set, two pairs documented as half-siblings were consistently classified as avuncular by our across-chromosome phasing framework (*P* (HS) = 7 *×* 10^*−*5^ and *P* (HS) = 0.43). Rather than adjudicating label fidelity based on classifier output, an independent re-examination of the extended pedigree graph revealed that the first discordant pair (*P* = 7 *×*10^*−*5^) arose from cryptic two-sided relatedness across both parental lineages. As detailed in the Supplemental Material, this bilateral sharing falsely satisfied our first-cousin triangulation criteria while generating an avuncular-like alternating gametic structure, indicating that this discrepancy represents label noise in the reference set rather than algorithmic misclassification. For the second discordant pair (*P* = 0.43), no conflicting genealogical links or anomalous sharing topologies were detected in the pedigree graph; given that its posterior probability lies close to the classification boundary, the most plausible explanation is reduced across-chromosome phasing fidelity in one or both individuals due to a sparse local haplotypic neighborhood.

### 3.4 Comparison to other methods

We benchmarked our approach against two distinct classes of methods: network-based algorithms that rely on mutual relatives, and pairwise methods that evaluate segment sharing exclusively between the two target individuals.

For network-based inference, CREST (9) requires a minimum autosomal genome coverage rate (*C ≥* 0.025) across shared mutual relatives to separate second-degree classes reliably. In our HCPI validation set, only 92% of pairs satisfied this criterion; for the remaining 8% of isolated pairs lacking sufficient mutual relative coverage, CREST defaults to an uninformative prior. This requirement limits the operational recall of CREST in biobank cohorts containing sparse or isolated family clusters, an operational limitation our pairwise framework directly overcomes.

For pairwise inference, PONDEROSA (14) evaluates segment allocation across homologs using an within-chromosomal haplotype score, *h*, leveraging the biological expectation that half-siblings inherit segments exclusively on a single parental homolog (Figure 4). While this formulation is effective in datasets with near-perfect phasing, its discriminative accuracy degrades in the presence of phase errors that downstream error-correction heuristics fail to detect. To evaluate this approach on our cohort, we implemented the phase error handling, segment detection protocol, and parameters described in PONDEROSA (https://github.com/williamscole/PONDEROSA) under the standard recommended settings for `phasedibd` (17): merging gaps *<* 1.0 cM and discarding fragments *<* 2.5 cM around phase errors. We then computed *h* across all 22 autosomes and trained the linear discriminant analysis (LDA) classifier. In our biobank dataset, long-range phase errors fragmented homologous blocks, causing 39 of the 99 true half-sibling pairs to be misclassified into the phase-error or avuncular partitions (sensitivity of 0.61) and 13 avuncular pairs to be misclassified as half-siblings, resulting in an overall accuracy of 0.90. By contrast, our across-chromosome phasing formulation models inter-chromosomal segregation directly, remaining robust to local within-chromosomal phase errors and achieving near-perfect discrimination without requiring external pedigree members.

### 3.5 Resolving Directionality in Avuncular Pairs

Beyond distinguishing avuncular from half-sibling relationships, the asymmetric transmission of gametes across generations allows our across-chromosome phasing framework to resolve generational directionality— specifically, identifying which individual is the avuncular and which is the niece/nephew. For an avuncular pair, the niece/nephew inherits paternal or maternal chromosomal segments that trace back to their grandparents through a single connecting parent. Consequently, all shared IBD segments in the niece/nephew reside exclusively on one parental chromosome, whereas the avuncular shares segments inherited across both of their own parental homologs with that same single transmitted homolog of the niece/nephew (Figure 4).

We operationalized this structural asymmetry by evaluating the entries of the 2 *×* 2 haploid sharing matrix 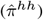 defined in Table 1. After ordering the matrix so that 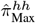 represents the cell with maximal across-chromosome segment overlap, the remaining sharing along the row and column of 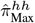 yields two directional values: 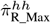 and 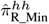 . In an avuncular pair, biological transmission dictates which of these two cells corresponds to true ancestral sharing: the avuncular’s second homolog sharing with the transmitted niece/nephew homolog should remain substantial 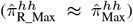, whereas the niece/nephew’s unshared, opposite-parent homolog captures only background haplotypic Noise 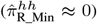. Which dimension (row or column) contains 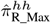 therefore indicates which individual carries the two transmitting homologs, directly resolving who is the avuncular in the pair.

When genealogical roles are known from reference pedigree links, the transmitting individual is established by identifying which member is the avuncular relative. This designates the specific matrix cell expected to hold 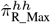. Therefore, under correct generational assignment, the two expected values for 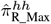 and 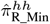 satisfy:

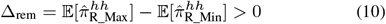

We validated this directionality metric across all 444 candidate avuncular pairs in our HCPI reference set where generational roles were independently determined via reconstructed third-party pedigree links (i.e., whether individual 1 was an avuncular via a sibling-parent link or individual 2 was an avuncular via a parent-sibling link). Consistent with theoretical predictions, designating the expected 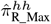 by the documented reference role clearly polarizes the pair: Δ_rem_ follow a positive distributions (Supplementary Figure S6). Among the 444 evaluated reference avuncular pairs, only a single pair exhibited an inverted sign (Δ_rem_ *<* 0), yielding an empirical directionality assignment accuracy of 99.77% (443*/*444 pairs) in resolved avuncular relationships. These results confirm that across-chromosome phasing not only discriminates degree-2 classes without pedigrees, but also polarizes parent-offspring generational order within avuncular dyads.

## 4 Application to Phasing

Beyond identification of relative types, inferred half-sibling and avuncular relationships can be leveraged as constraints to improve both within- and across-chromosome phasing. For half-sibling pairs, all shared IBD segments derive from a single common parent, providing unambiguous information about parental haplotype origin and thus serving as effective anchors for phase assignment across the genome. For the avuncular in avuncular pairs, shared segments cannot generally be assigned to a specific parental lineage without additional information, because they may derive from either grandparental haplotype carried by the avuncular relative. However, these segments are informative for phasing the niece or nephew, as they must have been transmitted through the intervening parent, thereby constraining haplotype assignment in that individual. When incorporated into the reference panel of a phasing algorithm, such relationships increase the consistency and connectivity of inferred haplotypes across individuals and can improve overall phasing accuracy.

We incorporated these relatives into the across-chromosome phasing framework described in Sapin et al. (11) by explicitly including second-degree relatives as across-chromosome phase anchors. Following the definition established in Sapin et al. (11), across-chromosome phasing accuracy was evaluated as the percentage of the genome where the chromosome-specific haplotypes were correctly assigned to a consistent parental origin. For HCPI nieces/nephews where parental geno-types permitted direct validation (whereas for most half-sibling pairs no parental genotype is available to assess phasing accuracy), this relationship-informed constraint markedly enhanced performance. Specifically, the introduction of second-degree anchors increased the mean across-chromosome phasing accuracy from 84.04% to 92.76%, while the median accuracy improved from 86.56% to 93.40%. These results demonstrate that subtyping second-degree relatives provides a robust, genotype-only feedback loop for refining haplotypic reconstruction in biobank-scale datasets.

## 5 Data and Code Availability

The underlying genomic and genealogical data used in this study were obtained from the UK Biobank Resource, which is a restricted-access dataset. Access to individual-level genotype arrays and phenotypic records can be requested by bona fide researchers directly through the UK Biobank Access Management System.

All custom source code developed for this study—including the multithreaded C++ implementation for pairwise across-chromosome IBD feature extraction (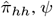, and homolog assignment) and the R pipelines for two-dimensional Gaussian Mixture Model (GMM) fitting, out-of-fold cross-validation, and posterior probability estimation—is publicly available in our GitHub repositories: https://github.com/esapin/Across-Chromosome-Phasing and https://github.com/esapin/HSNA. Step-by-step reproduction instructions, sample input formats, and parameter configurations are documented in the repository’s `README.md`.

## 6 Conclusion and Future Work

This study demonstrates that a multivariate Gaussian mixture model applied to haplotype-level sharing features derived from acrosschromosome phasing can reliably distinguish half-sibling from avuncular (niece/nephew–uncle/aunt) pairs using genotype data alone, providing a scalable alternative to approaches that rely on explicit pedigree structures or phenotypic data. By exploiting the asymmetric inheritance of parental homologs, the method resolves second-degree relationship classes that remain indistinguishable using aggregate diploid SNP metrics. In the UK Biobank, our supervised framework achieved 97.98% specificity and 100% sensitivity on the high-confidence reference set, while polarizing generational roles within avuncular pairs (identifying avuncular versus niece/nephew) with 99.77% empirical accuracy.

Compared to existing relatedness inference methodologies, this approach addresses critical operational bottlenecks in biobank-scale cohorts. Multi-way pedigree reconstruction algorithms such as CREST (9) rely on shared mutual relatives, leaving isolated pairs lacking extensive kindred networks uninformative. Conversely, pairwise methods that leverage within-chromosome haplotype sharing, such as PONDEROSA (14), can degrade substantially when phase errors break phase continuity along individual autosomes. By modeling gametic transmission across chromosomes rather than within chromosomes, our framework remains robust to local phase errors and operates effectively on isolated dyads without requiring collateral relatives or in-sample labeled training pedigrees.

We also demonstrate that resolving second-degree relationships establishes an informative feedback loop for genomic reconstruction. Because half-siblings inherit all shared segments from a single shared parent, they provide unambiguous parental haplotype constraints across the entire autosome; similarly, segments shared between an avuncular and niece/nephew constrain haplotype phasing in the niece/nephew. Incorporating these inferred pairs as phase anchors directly enhanced across-chromosome phasing fidelity, increasing the mean phasing accuracy from 84.04% to 92.76% in validated offspring.

An important limitation of the current work is its focus on the two dominant second-degree classes in adult cohorts. Our framework does not explicitly model rarer degree-2 configurations such as double first cousins and grandparent–grandchild (GP–GC) pairs. While double first cousins are readily identified and removed via non-zero 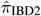 sharing 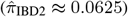, true GP–GC pairs share the same uniparental gametic transmission profile as avuncular pairs and would cluster with them in our feature space. Although demographic age constraints (enrollment ages 40– 69) render GP–GC pairs negligible in the UK Biobank, cohorts spanning multiple generations would require joint modeling of age distributions or genealogical depth to separate avuncular from grandparental lineages.

A natural extension for future work is generalizing this haplotype-aware paradigm to higher-degree relationships. In third-degree pairs sharing approximately 12.5% of the genome, separating first cousins from half-avuncular pairs presents an analogous symmetry challenge, as both involve first-degree relationships among the connecting parents. By reconstructing the four grand-parental haplotypic origins across chromosomes, it may be possible to differentiate single-lineage transmission from alternating grand-parental segregation. Additionally, extending across-chromosome phasing frameworks into multi-ancestry cohorts will be essential to evaluate the impacts of recombination landscape diversity, endogamy, and varying background kinship on classification power.

In summary, across-chromosome phasing provides a principled, genotype-only mechanism to untangle second-degree relationships and resolve parent-offspring generational order. By removing the dependence on observed pedigrees or dense family clusters, this methodology improves relationship annotation, sample quality control, haplotype phasing, and downstream variance decomposition across large-scale genomic biobanks.

## 7 Declaration of Generative AI

During the preparation of this work, the authors used Gemini (November 2026 version) to improve language clarity, structure paragraphs for the revised manuscript text, and ensure technical syntax alignment within the manual bibliography section. The AI-assisted tools were used under the direct monitoring and supervision of the authors, who take full responsibility for the content of the published article.

