## Supplement for "Near-perfect discrimination of half-sibling and avuncular pairs using genomic data without pedigrees"

### 1 Ground Truth Identification

As described in the main text, we define a High-Confidence Pedigree-Inferred (HCPI) validation set to serve as ground truth for avuncular pairs was established using direct pedigree links within the dataset. However, the HCPI validation set for half-sibling pairs required a multi-faceted approach involving three distinct methods: common parentage, shared first cousins, and unrelated first cousins.

#### 1.1 Avuncular Pairs

The HCPI validation set for avuncular pairs was constructed using genotyped trios where an individual is a sibling to one member of the target pair and the parent of the other. Parent-offspring and sibling pairs were identified using a threshold of $\hat{\pi} > 0.33$. A threshold of $\hat{\pi}_{\mathrm{IBD2}} > 0.1$ was then applied to differentiate full siblings from parent-offspring pairs (where $\hat{\pi}_{\mathrm{IBD2}} \approx 0$).

#### 1.2 Half-Sibling Identification via Shared Pedigree

Direct identification of half-siblings through a common genotyped parent yielded only three pairs. To expand the HCPI validation set, we leveraged the presence of first cousins. This approach necessitates precise thresholds on $\hat{\pi}$ and $\hat{\pi}_{\mathrm{IBD2}}$ to avoid misclassification due to complex pedigree structures.

#### 1.3 Validation via First Cousin Triangulation

As detailed in the main text, two individuals sharing a first cousin must be half-siblings, provided the cousin is related through the shared parental lineage. Initially, relatives were selected as first cousins based on $0.115 < \hat{\pi} < 0.14$. However, this threshold alone proved insufficient, occasionally leading to the misidentification of first cousins due to double-lineage effects.

For example, a pair $(\mathrm{Gao}_1, \mathrm{Gao}_2)$ with $\hat{\pi} = 0.236$ was initially misclassified as half-siblings because a relative, $W$, appeared to be a first cousin to both (see Figure 1). While $W$ shared a $\hat{\pi}$ of $0.128$ with $Gao_1$ and $0.134$ with $Gao_2$, the $\hat{\pi}_{\mathrm{IBD2}}$ value with $Gao_2$ was $0.0063$. This positive $\hat{\pi}_{\mathrm{IBD2}}$ suggests a double-lineage link between $Gao_2$ and $W$, which can be explained by the relatedness situation described in Figure 1. In this configuration, both parents $P_2$ and $P_7$ of $Gao_2$ are first cousins of $W$. Such a structure is incompatible with a standard half-sibling relationship between $Gao_1$ and $Gao_2$. If they were half-siblings, the relatedness of $W$ to $Gao_2$ coming the parent in common between $Gao_1$ and $Gao_2$ would be expected to remain within the same range as the relatedness of $W$ to $Gao_1$ and additionaly the relatedness of $W$ to $Gao_2$ coming from the other parent of $Gao_2$ would raise the relatedness to a value of $\hat{\pi}$ away from the degree 3 range; therefore the observed values instead pointed to an avuncular relationship.

Further validation of the avuncular status of $(Gao_1, Gao_2)$ was provided by a relative $Z$. $Z$ shared a $\hat{\pi}$ of $0.052$ with $Gao_1$ but only $0.024$ with $Gao_2$ (approximately half the value). Given that $\hat{\pi}_{\mathrm{IBD2}} < 0$ for both, which excludes double-lineage relatedness, this pattern suggests that $Z$ could a first cousin once removed to $Gao_1$ and a first cousin twice removed to $Gao_2$. This differential distance confirms the avuncular relationship (Degree 3 vs. Degree 4 distance from $Z$). While other interpretations of these values are mathematically possible—such as $Z$ being a great-grand-avuncular relative of $Gao_1$—the restricted age range of individuals in the UK Biobank makes such an intergenerational hypothesis highly improbable.

#### 1.4 First Cousin Inclusion Constraints

To mitigate uncertainty, we established a formal control: for every potential first cousin ($PFC$) of either individual in a pair where $0.0975 < \hat{\pi} < 0.1875$, the value with the other individual must satisfy one of two conditions:

- **Unrelated through the unshared parent**: $\hat{\pi} < 0.025$.
- **Related through the shared parent**: $\hat{\pi} > 0.0975$.

Any Degree 2 pair containing an individual with a $PFC$ that fails to meet either of these constraints was excluded from the HCPI validation set to ensure maximum specificity.

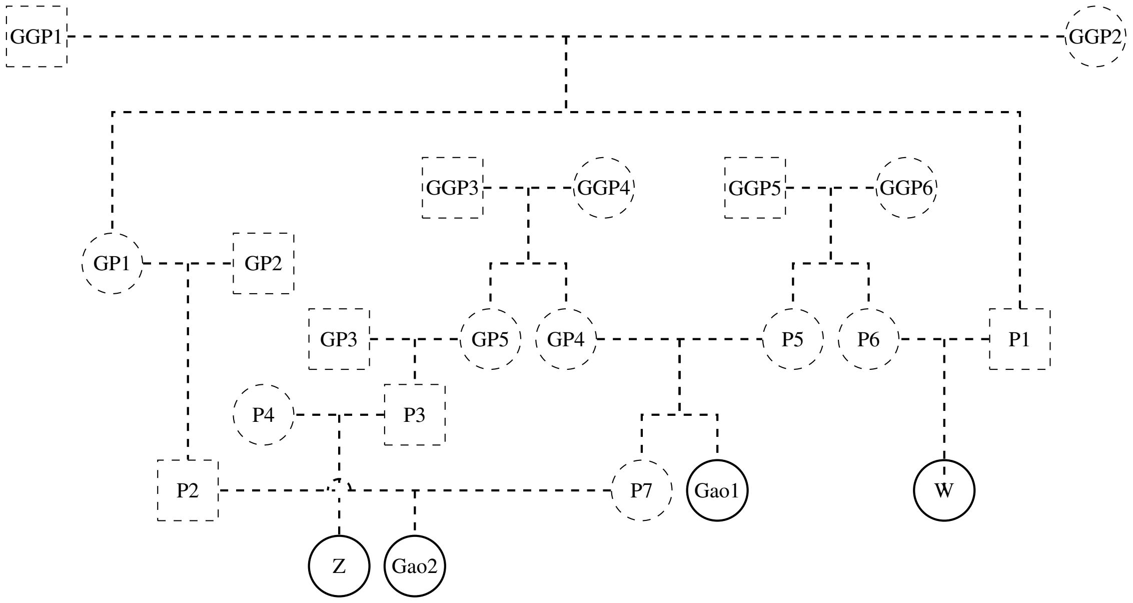


Figure 1: Example pedigree with a two-sided first cousin once removed

### 2 Posterior Distribution and Boundary Resolution

While the main text displays the distribution of posterior probabilities on a linear scale to illustrate population-level proportions, Figure 2 presents the same posterior probabilities, $P(\text{half-sibling})$, on a $\log_{10}$ frequency scale. Visualizing frequencies logarithmically highlights the low-density boundary region ($0.1 < P(\text{H-S}) < 0.9$) that is otherwise obscured by the extreme mass at the polar extremes ($P \approx 0$ and $P \approx 1$).

This logarithmic view highlights two critical aspects of our classifier's performance:

1. **High Specificity for Avuncular Relatives:** Across all $n = 444$ ground-truth avuncular pairs, none cross the $P(\text{H-S}) \geq 0.5$ decision boundary, with the highest observed avuncular pair reaching $P = 0.23$.
2. **Granular Characterization of Boundary Cases:** The log scale directly resolves the two misclassified ground-truth half-sibling pairs ($P \approx 0.42$ and $P < 0.05$). Furthermore, for the unlabeled biobank candidate cohort ($n = 8,053$), the frequency density across the intermediate uncertainty region ($0.35 \leq P \leq 0.65$) remains over two orders of magnitude lower than at the polar boundaries, demonstrating that ambiguous classifications represent a very small fraction of candidate second-degree pairs.

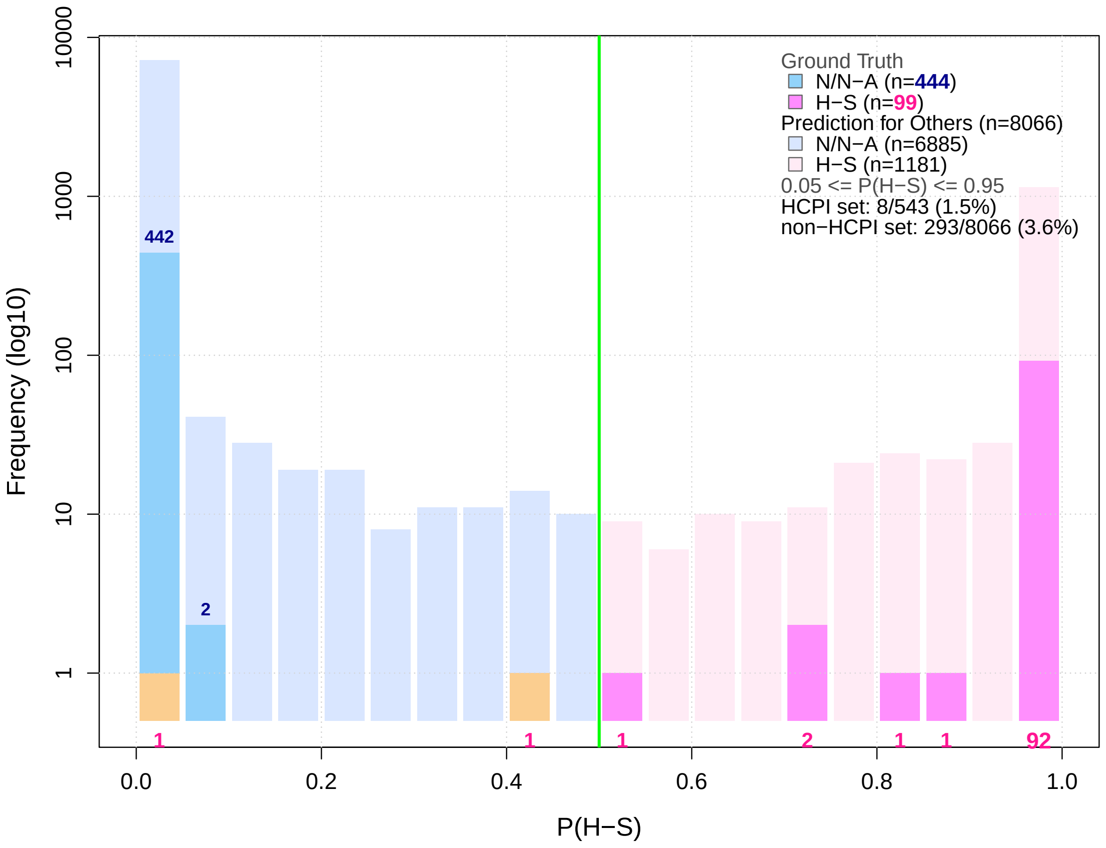


Figure 2: Distribution of the posterior probability $P(\text{half-sibling})$ on a log scale for HCPI set of pairs and unlabeled second-degree relative candidates. The solid green line indicates the decision threshold for half-sibling identification ($P(\text{half-sibling}) > 0.5$). Black numbers indicate number of HCPI avuncular relatives and red numbers indicate the number of HCPI half-siblings.

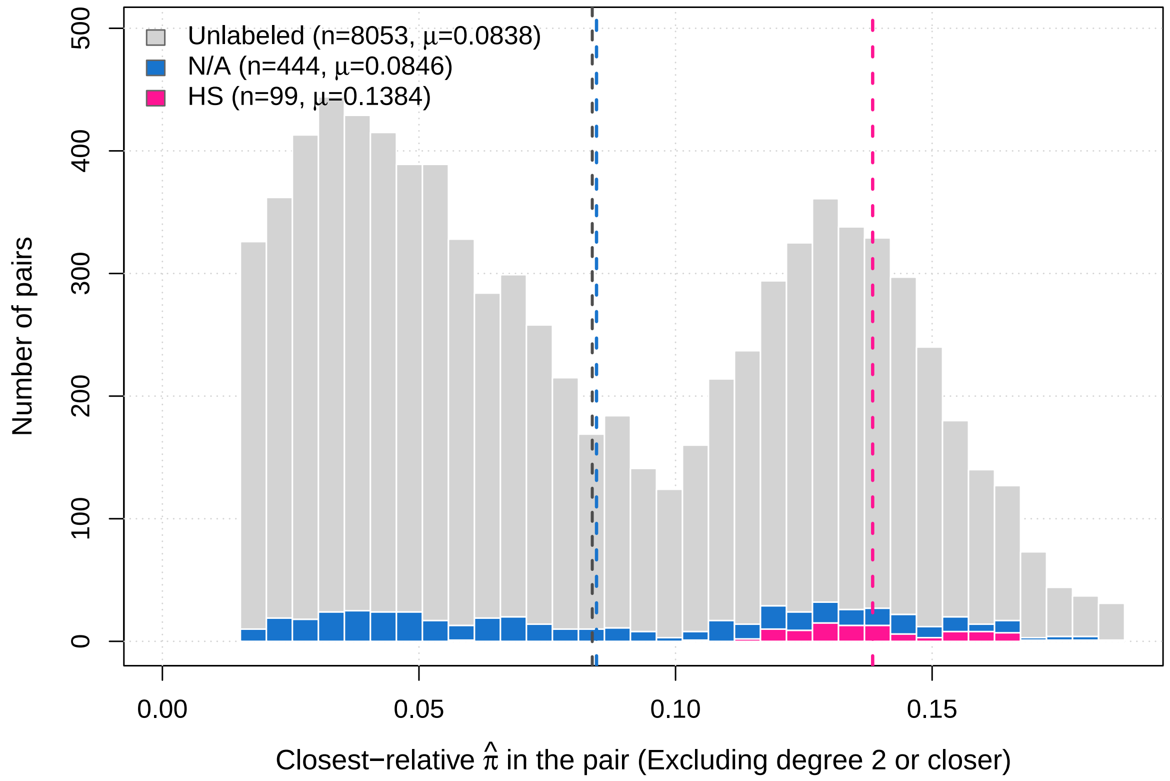


Figure 3: **Distribution of external relatedness across cohorts.** Closest external relatedness ($\hat{\pi} < 0.1875$) for unlabeled pairs ($n = 8{,}053$, grey), reference avuncular pairs ($n = 444$, blue), and reference half-siblings ($n = 99$, pink), with dashed lines indicating group means[cite: 15]. The unlabeled cohort closely matches the avuncular reference distribution.

### 3 External Relatedness Distributions

To quantify the familial background across cohort partitions, we evaluated empirical relatedness ($\hat{\pi}$) to each target pair's closest external relative, excluding first- and second-degree links ($\hat{\pi} < 0.1875$; Figure 3). The resulting histograms reveal distinct genealogical sampling profiles across the reference and unlabeled subsets. For ground-truth avuncular pairs, the closest external relatedness is centered at a mean of 0.0846, reflecting moderate familial embedding within the biobank. In contrast, ground-truth half-sibling pairs exhibit a shifted distribution with a substantially higher mean of 0.1384, as expected from their ascertainment through collateral first-cousin triangulation. Crucially, the distribution across the 8, 053 unlabeled candidate pairs closely mirrors that of the avuncular reference group, exhibiting a mean external relatedness of 0.0838 and demonstrating that the unlabeled biobank cohort primarily consists of genealogically isolated pairs rather than extended multi-generational kindreds.

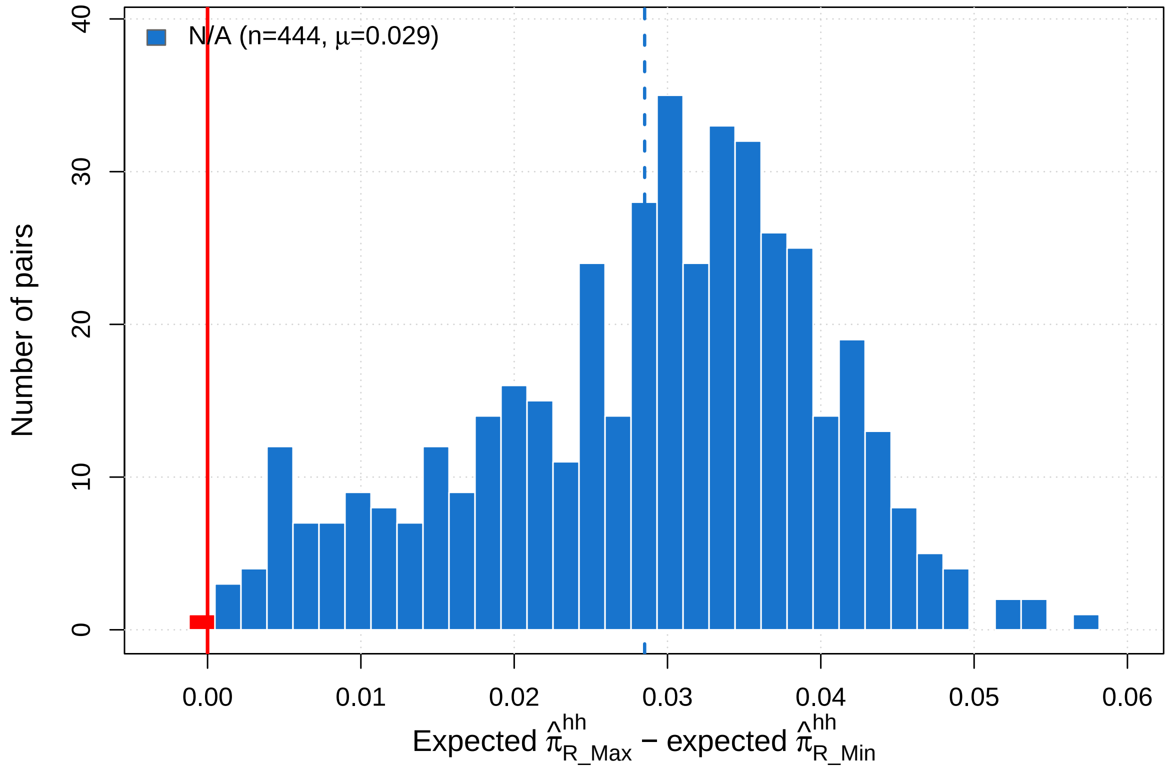


Figure 4: **Directionality metric $\Delta_{\mathbf{rem}}$ in reference avuncular pairs.** Distribution of $\Delta_{\text{rem}} = \mathbb{E}[\hat{\pi}^{hh}_{\text{R\_Max}}] - \mathbb{E}[\hat{\pi}^{hh}_{\text{R\_Min}}]$ ($n = 444$, mean $\mu = 0.029$ dashed line). Values $> 0$ correctly polarize generational directionality ($443/444$ pairs, $99.77\%$ accuracy).

### 4 Resolving Directionality in Avuncular Pairs

Because an avuncular relative transmits segments across both parental homologues to a single homologue of the niece or nephew, generational directionality can be resolved by comparing the two directional sharing sums along the row and column containing $\hat{\pi}^{hh}_{\text{Max}}$ (Figure 4). In reference avuncular pairs where genealogical roles were determined by independent pedigree links, designating the transmitting relative as the avuncular yields a directional difference $\Delta_{\text{rem}} = \mathbb{E}[\hat{\pi}^{hh}_{\text{R\_Max}}] - \mathbb{E}[\hat{\pi}^{hh}_{\text{R\_Min}}]$ that is positive across 443 of 444 pairs (mean $\mu = 0.029$; empirical polarization accuracy of $99.77\%$).

### 5 PONDEROSA Classification Benchmark and Posterior Distribution

To benchmark our across-chromosome phasing approach against pairwise within-chromosome haplotype sharing methods, we implemented the segment detection protocol, phase-error handling heuristics, and classification parameters described in PONDEROSA (Williams et al., 2025; see Main Text) using `phasedibd` (Freyman et al., 2021). For each second-degree candidate pair, identity-by-descent (IBD) segments were detected under standard recommended settings (merging adjacent segments separated by gaps $< 1.0$ cM and discarding fragments $< 2.5$ cM proximal to detected phase errors). The within-chromosome haplotype score $h$ was aggregated across all 22 autosomes, and a linear discriminant analysis (LDA) classifier was trained to separate half-sibling (H-S) from avuncular (N/N-A) relationships.

Figure 5 displays the resulting distribution of posterior probabilities, $P(\text{H-S})$, on a $\log_{10}$ frequency scale across both the High-Confidence Pedigree-Inferred (HCPI) validation cohort ($n = 543$) and the unlabeled candidate second-degree cohort ($n = 8,058$). The vertical green line represents the empirical decision boundary at $P(\text{H-S}) \approx 0.43$, calibrated to maximize overall classification accuracy across HCPI reference pairs rather than using the default equal-cost LDA threshold of 0.5.

Unlike the sharp, polar separation achieved by our across-chromosome GMM framework (where posterior densities in the intermediate uncertainty interval $0.1 < P(\text{H-S}) < 0.9$ remain over two orders of magnitude lower than the polar extremes), the PONDEROSA baseline exhibits substantial mass throughout the intermediate region ($0.2 \leq P(\text{H-S}) \leq 0.7$). In population-scale cohorts such as the UK Biobank, long-range phase errors along individual autosomes frequently truncate and fragment homologous shared blocks. This fragmentation causes $n = 39$ of the 99 HCPI half-sibling pairs to receive attenuated haplotype scores, shifting their posterior probabilities below the decision threshold into the avuncular partition (yielding a half-sibling sensitivity of only $60.61\%$). Similarly, $n = 13$ HCPI avuncular pairs are misclassified as half-siblings, resulting in an overall HCPI validation accuracy of $90.42\%$ (491/543 pairs correctly classified).

When applied to the unlabeled candidate cohort ($n = 8,058$), the classifier assigns 6,871 pairs as avuncular and 1,187 pairs as half-siblings. However, the substantial density spanning the classification boundary underscores the sensitivity of within-chromosome scoring to unresolved phase errors, demonstrating why aggregating gametic transmission profiles across chromosomes provides far greater discriminative stability in biobank cohorts.

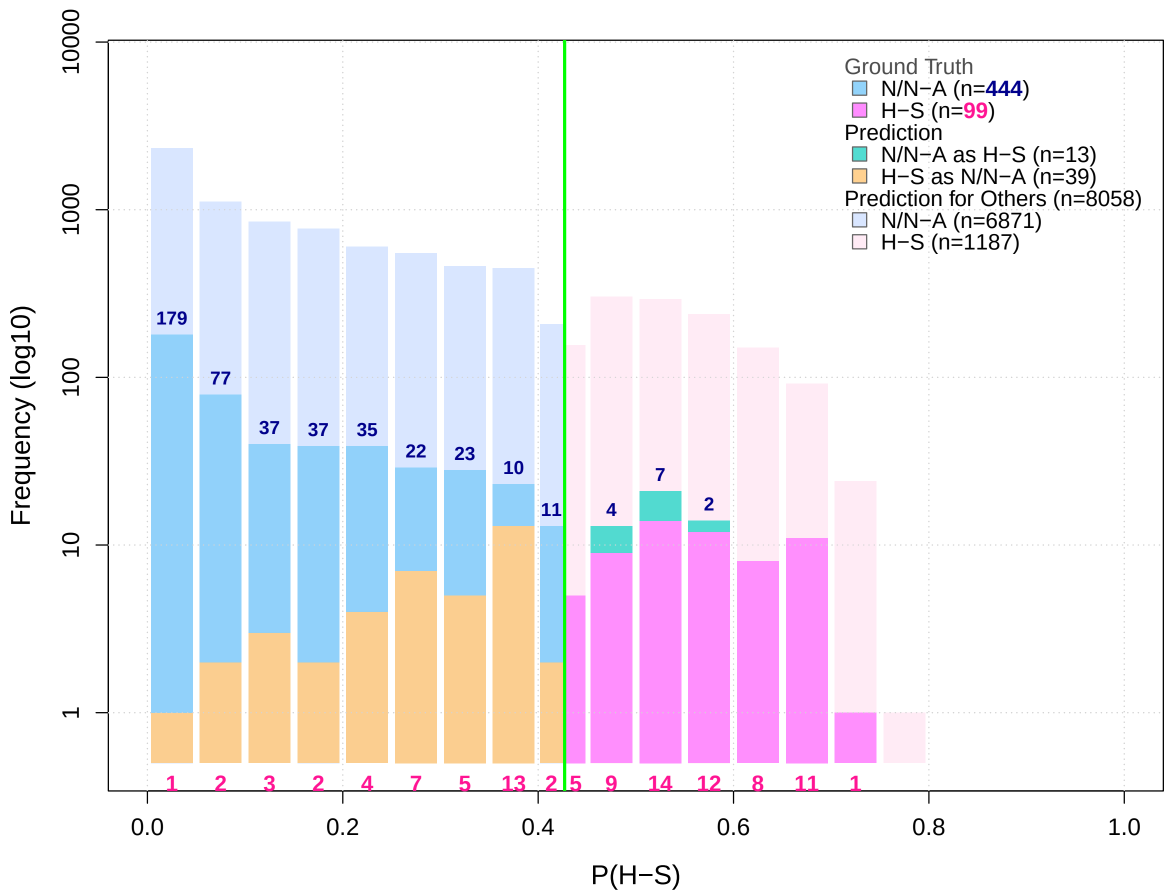


Figure 5: **Distribution of posterior probabilities $P(\textbf{H-S})$ under the PONDEROSA within-chromosome framework on a $\log_{10}$ frequency scale.** The histogram shows classification scores for HCPI validation pairs ($n = 444$ avuncular [blue] and $n = 99$ half-sibling [magenta]) alongside predictions across the unlabeled cohort ($n = 8,058$; shaded bars). Misclassifications within the HCPI reference cohort are highlighted in orange (HCPI half-siblings misclassified as avuncular, $n = 39$) and cyan (HCPI avuncular pairs misclassified as half-siblings, $n = 13$). Dark blue numbers indicate counts of correctly classified avuncular pairs per bin; magenta numbers below the horizontal axis indicate counts of half-sibling pairs. The solid green line indicates the operational decision boundary at $P(\text{H-S}) \approx 0.43$, chosen to maximize empirical accuracy on the HCPI validation set rather than using the equal-cost LDA threshold of 0.5. Overall classification accuracy across the HCPI validation set was 90.42% (491/543 pairs), with half-sibling sensitivity reduced to 60.61% (60/99 pairs) as a consequence of long-range phase errors.
